# Head Mesoderm Tissue Growth, Dynamics and Neural Crest Cell Migration

**DOI:** 10.1101/738542

**Authors:** Mary Cathleen McKinney, Rebecca McLennan, Rasa Giniunaite, Ruth E. Baker, Philip K. Maini, Hans G. Othmer, Paul M Kulesa

## Abstract

Vertebrate head morphogenesis involves orchestrated cell growth and tissue movements of the mesoderm and neural crest to form the distinct craniofacial pattern. To better understand structural birth defects, it is important that we learn how these processes are controlled. Here, we examine this question during chick head morphogenesis using time-lapse imaging, computational modeling, and experiment. We find that head mesodermal cells are inherently dynamic in culture and alter cell behaviors in the presence of either ectoderm or neural crest cells. Mesodermal cells in vivo display large-scale whirling motions that rapidly transition to lateral, directed movements after neural crest cells emerge. Computer model simulations predict distinct changes in neural crest migration as the spatio-temporal growth profile of the mesoderm is varied. BrdU-labeling and photoconversion combined with cell density measurements then reveal non-uniform mesoderm growth in space and time. Chemical inhibition of head mesoderm proliferation or ablation of premigratory neural crest alters mesoderm growth and neural crest migration, implying a dynamic feedback between tissue growth and neural crest cell signaling to confer robustness to the system.

**Summary Statement:** Dynamic feedback between tissue growth and neural crest cell migration ensures robust neural crest stream formation and head morphogenesis shown by time-lapse microscopy, mathematical modeling and embryo perturbations.

## INTRODUCTION

Neural crest cell migration is crucial to vertebrate head development since multipotent cells must reach and contribute to morphogenesis of the face and branchial arches. Signals within the hindbrain control neural crest cell exit locations and sculpt cells into discrete streams. To reach the periphery after exiting the dorsal neural tube, cranial neural crest cells travel through dense extracellular matrix and mesoderm, subjacent to the ectoderm (Tosney, 1982; Noden, 1988). Tissue transplantations, together with viral and fluorescent dye marking of the cranial neural crest and paraxial mesoderm have revealed that both cell types are migratory and follow stereotypical pathways into the branchial arches (Kontges and Lumsden, 1996; Hacker and Guthrie, 1998; Evans and Noden, 2006). However, these static analyses have not been able to tease out the cell dynamics and the relationship between cell proliferation, tissue growth and cell-cell interactions that contribute to the complexity of head morphogenesis. This is primarily due to the lack of a detailed in vivo analysis of head mesoderm and ectoderm dynamics in space and time, and limitations with in vivo imaging of neural crest-mesodermal cell interactions.

A better understanding of the dynamics and interplay between the neural crest and tissues through which cells travel would shed light on the mechanistic basis of collective cell migration for which neural crest cells are an exemplary model (Noden, 1988; Kontges and Lumsden, 1996; Noden and Trainor, 2005; McLennan et al., 2015; McKinney et al., 2016). Detailed experiments would confirm the robustness of the stereotypic neural crest stream shape and length over the course of early head development under the constraints of a growing microenvironment. In the absence of dynamic data, it is possible to speculate on a wide variety of tissue growth profiles for the head mesoderm and its impact on collective neural crest cell migration. In previous work, we quantified a curvilinear measurement of the distance from the chick neural tube dorsal midline along the dorsolateral cranial neural crest cell migratory pathway, to the tip of branchial arch 2 (ba2) throughout early developmental stages (McLennan et al., 2012; McLennan et al., 2017). This simple length measurement revealed a logistic growth profile of the total neural crest cell migratory domain over time (McLennan et al., 2017). However, in the absence of more detailed analyses, it has remained unclear whether head mesoderm growth is uniform or non-uniform in space during neural crest cell migration through this growing tissue.

In this study, we investigate the spatial and temporal dynamics of individual mesoderm and ectoderm cells, and their interplay with migrating neural crest cells during chick head morphogenesis. We computationally explore domains of different non-uniform in space mesoderm growth profiles and converge on a subset of possible outcomes that result in a continuous neural crest cell migratory stream. We examine and quantify the inherent migratory properties of mesoderm, either isolated from, or in the presence of neural crest cells, as a baseline for in vivo comparison. We then track and observe head ectoderm, mesoderm and neural crest cell behaviors simultaneously in Tg(hUBC: H2BCerulean-2A-Dendra2) whole quail embryo explants. We analyze the trajectories of individual mesodermal and neural crest cells in the same embryo, quantify the density fluctuations in the tissue, and selectively mark and measure changes in three-dimensional cell volume subregions within the ectoderm and mesoderm using photoconversion of small cell volumes in unprecedented detail. We image BrdU and methyl green labeling in cleared whole chick embryos to distinguish proliferative subregions within head mesoderm throughout successive chick developmental stages during neural crest cell migration. To address how changes in either the presence of neural crest cells or mesodermal cell proliferation affect cell dynamics and branchial arch size, we ablate a subregion of premigratory neural crest cells and measure subsequent tissue growth. Separately, we apply aphidicolin and hydroxyurea by focal microinjection into paraxial mesoderm to inhibit cell proliferation and compare the distribution of neural crest cells along the migratory domain with our computational model predictions. Together, our data offer the first detailed characterization of cell and tissue growth dynamics in the vertebrate head and a mechanistic basis for how neural crest cells must respond to migrate in a collective and directed manner.

## RESULTS

### Head mesoderm placed in culture is inherently dynamic and cell behaviors are influenced by co-culture with either neural crest cells or ectoderm

The growth of the vertebrate head is substantial during early development and migration of neural crest cells (Fig. 1A). To better understand the inherent migratory properties of individual cranial paraxial mesoderm cells, we visualized and quantified cell trajectories of Hoechst-labeled mesoderm isolated from the embryo lateral to rhombomere r4 at HH6-8 (Hamburger and Hamilton, 1951) (Fig. 1B). Cultured mesoderm moved rapidly to disperse throughout the culture dish with an average cell speed of approximately 64 um/hr and directionality (defined as distance from start to finish divided by the total distance traveled) of 0.22 (Fig. 1C,D). Mesoderm cell behaviors in culture resembled migrating neural crest cells with long, extended cellular processes that stretched between neighboring cells and open spaces of the culture dish (Movies 1,2).

**Figure 1:**
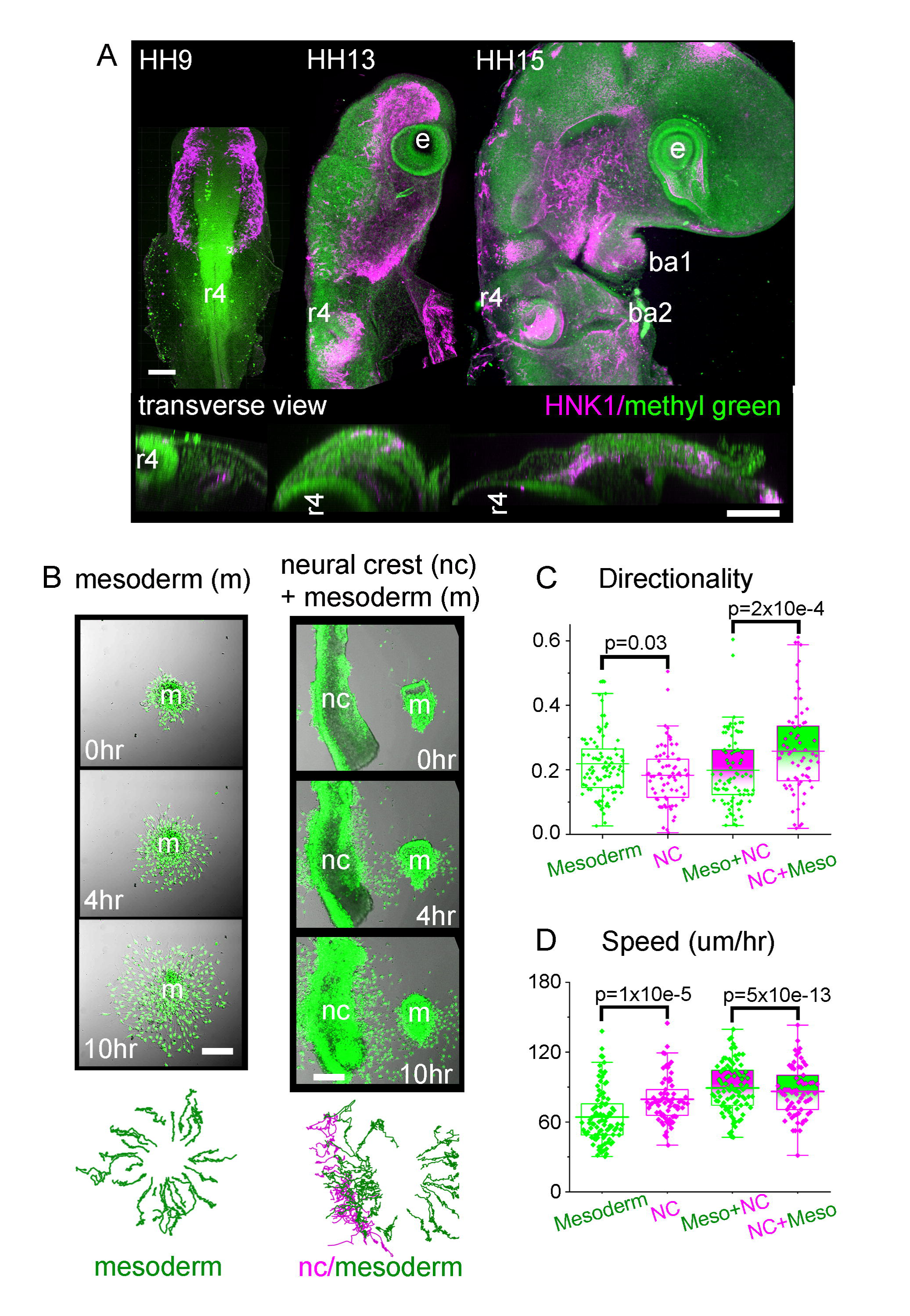
Growing chick embryo and mesodermal cell migration. (A) Similarly scaled chick embryos between stages HH9 and HH15 showing the growth of the head. All nuclei (green) and neural crest (magenta) illustrate the increase in number of cells and size of the embryo over time. Upper images: lateral view, scale bar 150um. Lower images: transverse view at the level of r4, scale bar 50um. (B) Column 1: Mesodermal cells isolated from HH6-8 embryos grown in vitro. Final panel are tracks of individual cells over 10 hours. Column 2: Neural tube from HH11 embryo co-cultured with mesoderm sample. Final panel are tracks of neural crest cells (magenta) and mesoderm (green). Scale bars 50um. (C) Directionality and (D) Speed of mesoderm only (green, 94cells), neural crest cells only (magenta, 73cells), mesoderm cells in the presence of neural crest (green markers with magenta box, 91cells) and neural crest in the presence of mesoderm (magenta markers with green box, 66cells).

Occasionally, we observed a few neural crest cells in the mesoderm explant (identified by HNK-1), but typically these cells were absent (Fig. S1). For comparison, cranial neural crest cells that exited from explanted neural tubes moved with an average speed of 79 um/hr and directionality of 0.19. While neural crest cells are statistically faster and cell trajectories are more circuitous than the mesoderm, exploratory behaviors and interaction of neighbors are similar. When we cultured mesoderm and neural tube explants together (Movie 3), both cell types increased their speed to 89 and 86 um/hr, respectively. However, only the mesoderm cell speed increased significantly such that there was no difference in speed between neural crest and mesodermal cells (Fig. 1C). The directionality for each cell type was not affected by the co-culture of both cell types, but neural crest cell directionality is still statistically different from that of mesoderm cells (Fig. 1D). Individual neural crest and mesoderm cells mixed freely, and some neural crest cells could be observed to infiltrate the explanted intact mesoderm (Movie 3). When we maintained the ectoderm with the intact mesoderm, we found that mesoderm frequently adhered to the sheet of ectoderm and traveled with this tissue (Movie 4).

Thus, chick head mesoderm is capable of dissociating and making individual cell movements, exploring the dish in a very similar manner to neural crest cells except when bound to ectoderm.

### In vivo time-lapse imaging reveals distinct collective motions of head mesoderm and ectoderm cell behaviors

Time-lapse imaging and analysis of head mesoderm and ectoderm cell behaviors in Tg(hUBC:H2BCerulean-2A-Dendra2) quail embryos (Huss and Lansford, 2017) revealed several interesting phenomena (Fig. 2A). First, as neural crest cells are emerging from the neural tube and beginning to migrate, the ectoderm can be seen to bend and change shape as the embryo grows (Fig. 2B and Movie 5). The large morphological changes to the embryo alter the presumptive path of the neural crest from at first being relatively straight to a curved path following the ectoderm deflection (Fig. 2B and Movie 5). Second, mesoderm and ectoderm cells adjacent to the dorsal hindbrain moved in a counter-clockwise whirling pattern, prior to the emergence of neural crest cells (HH9; Fig. 2C-E). The whirling pattern of cells had a diameter of approximately 125 um and was centered adjacent to rhombomere 3 (r3) (Fig. 2C-E). Mesoderm and ectoderm cells rotated together at 0.2 degrees per minute, suggesting that one revolution would take approximately 30 hrs to complete. The whirling cells were at least 50 um deep into the tissue but the maximum depth could not be determined due to the lack of light penetration into the tissue. Interestingly, the whirling behaviors of the mesoderm and ectoderm stopped coincident with the emergence of the r4 neural crest cells (Movie 6). Further, the pattern of mesoderm cell behaviors on the left-hand side of the embryo rotated in a clockwise manner (data not shown). The whirling effect was not observed in other locations or at later time points.

**Figure 2:**
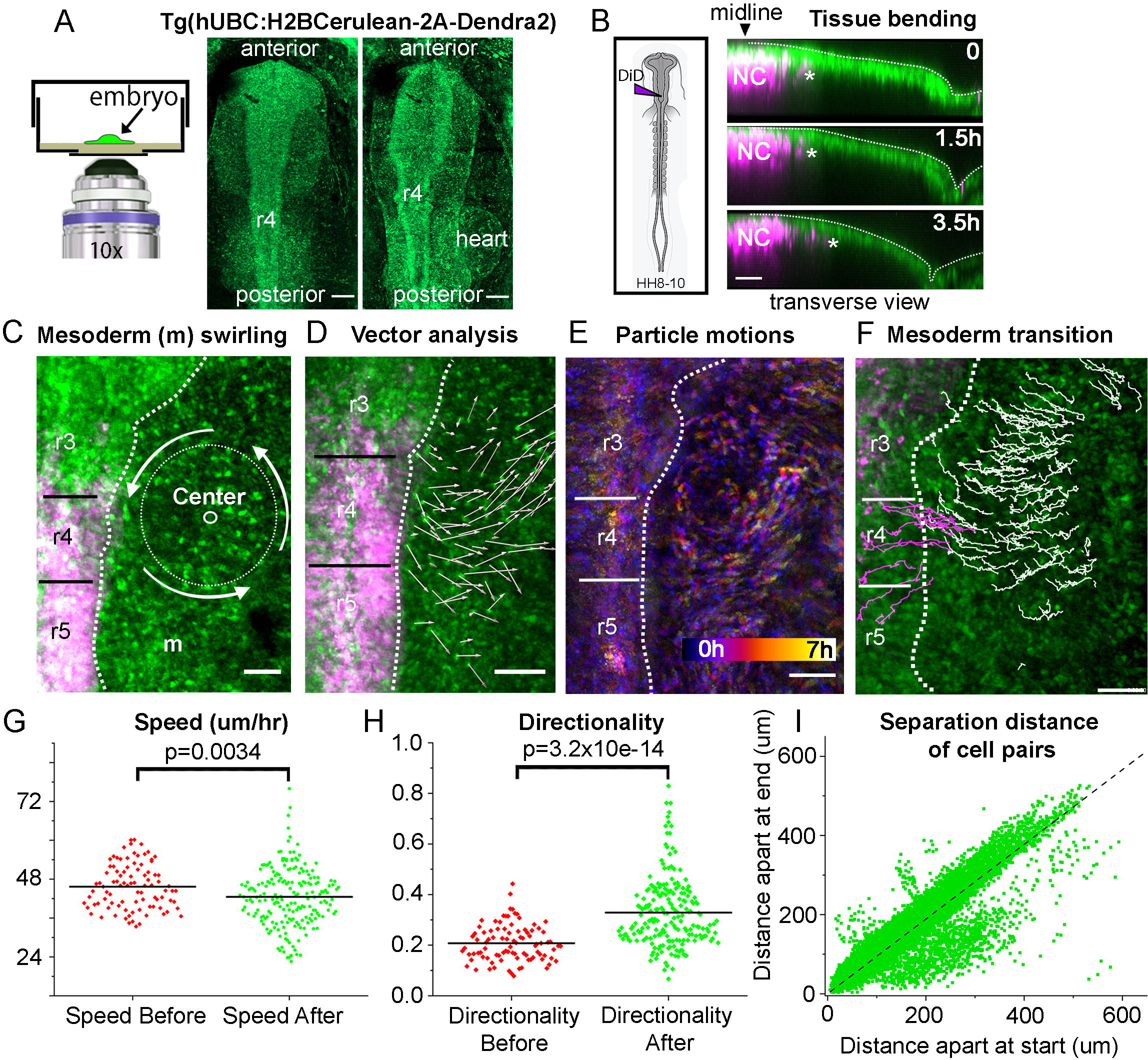
In vivo time-lapse imaging of head mesodermal cell behaviors. Representative images from 13 different time-lapse embryos. (A) Schematic of a typical embryo mounted with EC culture media. Images of Dendra2 (green) transgenic quail at HH9 and nine hours later. Scale bar 200um. (B) DiD labeling of premigratory neural crest in transgenic quail. Transverse view of r4 area over 3.5 hours in a time-lapse. Neural crest (magenta, leader marked with star) have just emerged from the neural tube while the surface ectoderm is changing shape. (C) Image of embryo at beginning of time-lapse with center of whirling motion and approximate size of circle marked. (D) Displacement of mesodermal and ectoderm cells (white arrows) over eight hours of imaging. (E) Color-coded time projected over a two-hour window with purple being time zero and yellow two hours later. The circular rotation of cells can be clearly seen. (F) Tracks of mesodermal cells (white) and neural crest (magenta) post neural crest emergence. Scale bars (B) – (F) are 50 um. Comparison of Speed (G) and Directionality (H) of mesodermal cells before and after the neural crest emerge from r4. (I) Comparison of the separation distance between any two cell pairs at the time of their first co-existence to their last observation, including pairs from different time-lapse movies. Straight line fit to data (dashed black).

### Mesoderm cell behaviors dramatically transition to directed, lateral trajectories towards the branchial arches after neural crest cells exit the dorsal neural tube

Third, we observed a dramatic change in mesoderm cell behaviors as neural crest cells exited the dorsal hindbrain in HH11 embryos. In contrast to large whirling behaviors near r3, mesoderm cells moved collectively in the proximal-to-distal direction (Fig. 2F). Cell tracking and analysis showed that mesoderm cells ‘before’ neural crest emergence moved at 46 um/hr on average with directionality of 0.21 (Fig. 2G,H). ‘After’ neural crest cells emerged from r4, mesoderm cells moved at a slightly slower speed of approximately 43 um/hr and with significantly higher directionality of 0.33—a 60% increase (Fig. 2G,H). Mesoderm cells were observed to move as a uniform collection of cells without trailing cells overtaking leaders (or leap-frogging) and without widespread cell rearrangements (Fig. 2F; Movies 6, 8). To examine this quantitatively, we measured the distance between random pairs of mesoderm cells over time (Fig. 2I). We find that neighboring mesoderm cells tended to remain neighbors, and cells far apart remained far apart as seen with cell pairs distributed along a straight line with slope approximately equal to 1 (Fig. 2I). We also found a smaller subpopulation of mesoderm cell pairs distributed below this line that shorten their distance between neighbors over time (Fig. 2I). Together, these data support our qualitative observations of the mesoderm moving as a continuous collection of cells growing and advancing in the proximal-to-distal direction to form the branchial arches.

### Neural crest cells migrate drastically different to neighboring mesodermal cells

When we co-labeled neural crest cells with DiD in Tg(H2BCerulean-2A-Dendra2) embryos and tracked neural crest and mesodermal cells over time (Fig. 3A,B), we discovered that neural crest cells moved through the mesoderm significantly faster (approximately 40% faster; 53 um/hr vs 38 um/hr for mesoderm) and with significantly higher directionality (0.4 vs 0.26 for mesoderm; Fig 3C,D). By the time that the neural crest cell migratory stream is fully defined adjacent to r4 (HH12), neural crest cells move at an even faster speed (97.7 um/hr, Movie 7). In contrast, individual ectodermal cells dorsal to the neural crest cell migratory pathway move slowly and are passed by the migrating neural crest cells (Movie 7). When we measured the distance between any two neural crest cells over time, we found that initially neighboring cells tended to remain neighbors throughout their migration (Fig. 3E). Analysis of individual neural crest and mesodermal cell interactions reveal that mesodermal behaviors varied in at least three ways (Fig. 3F, Movie 8). For example, a typical mesodermal cell positioned distal to the neural crest cell invasive front is eventually passed by rapidly moving neural crest cells (Fig. 3F, squares). For two hours, this mesodermal cell maintains a straight trajectory parallel to the passing, migrating neural crest cells, then changes direction to move posterior. This behavior contrasts the usual behavior seen in the neural crest cells themselves which generally stay in order and do not radically pass neighboring cells. Second, as migrating neural crest cells collide with mesodermal cells, the mesodermal cells are observed to match direction and move along with the neural crest stream as though the cells are pushed (Fig. 3F, circle). Measurements showed that mesodermal cells increased directionality after neural crest emergence (Fig. 2H). Third, some mesodermal cells located very distal to the emerging neural crest cell migratory stream may still be overtaken by the migrating neural crest cells (Fig. 3F, triangle).

**Figure 3:**
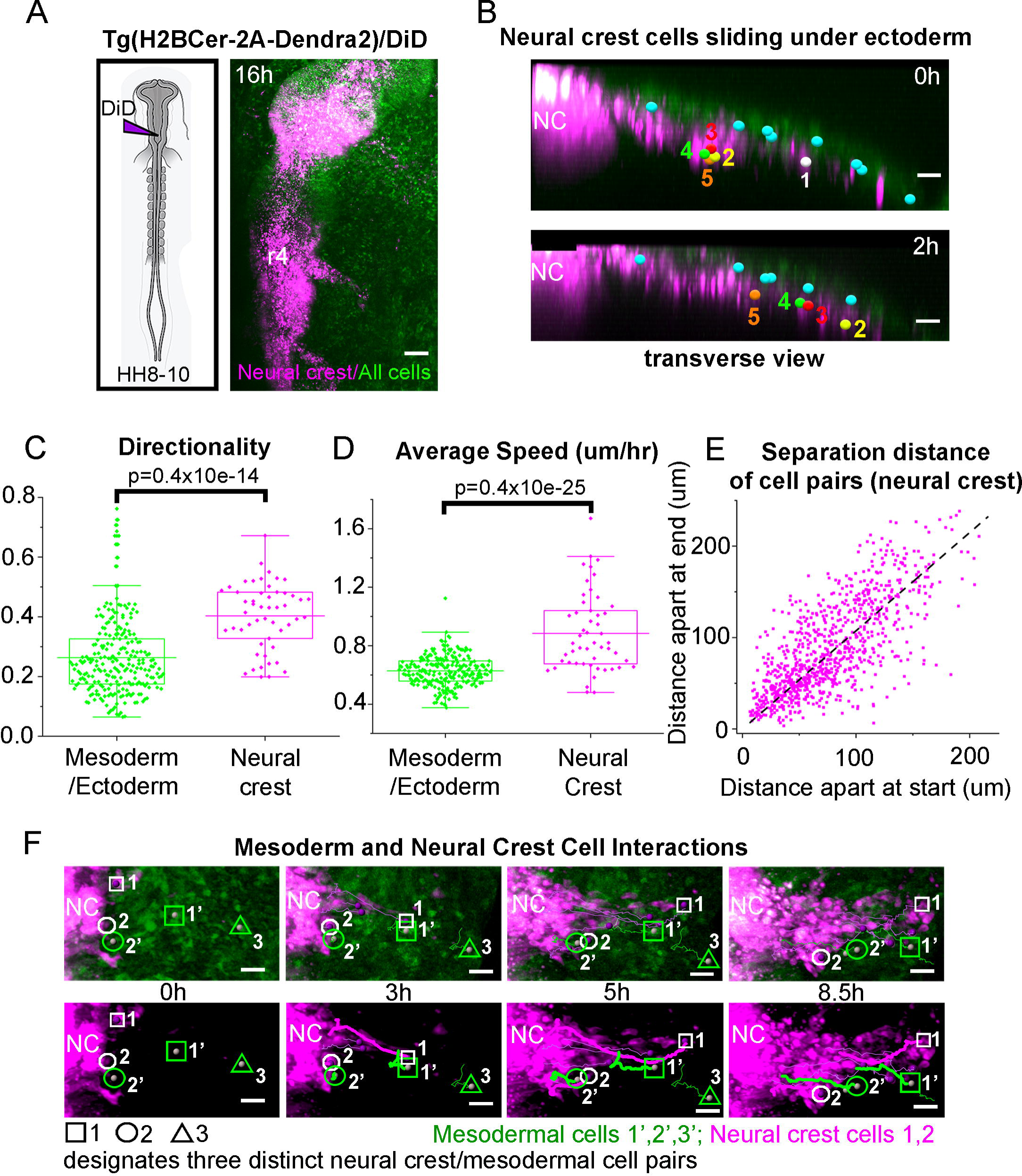
Neural crest cells are faster and more directed than the surrounding mesoderm. (A) DiD-injected embryo at +16hrs with neural crest (magenta) and Dendra2 (green). Scale bar 100um. (B) Transverse view at r4 at HH12. Individual ectoderm cells (cyan), neural crest cells and numbers showing the relative speed of neural crest compared to the neighboring tissue. Scale bar 20um. Comparison of Directionality (C) and Speed (D) of neural crest, mesoderm and ectodermal cells. (E) Comparison of separation distance from any two random neural crest cells at the time of their first co-existence to their last. Straight line fit to data (dashed black line). (F) Series of images from a typical time-lapse movie showing interaction between neural crest (magenta) and mesodermal cells (green) over 8.5 hrs. Circles (neural crest). Squares and Triangle (mesodermal cells). Scale bar 20um.

### Computer model simulations reveal that different tissue growth profiles on a 2D domain give rise to significantly distinct neural crest cell migratory stream distributions

To explore the effects of a set of distinct non-uniform domain growth profiles on the neural crest cell migratory pattern, we exploited the strengths of our computer model (Fig. 4). We considered six distinct non-uniform domain growth profiles in which the domain growth was twice as fast within either the proximal or distal subregions of the domain (Fig. 4A). Three out of the six growth profiles considered faster growing subregions within the distal half of the domain (Fig. 4A,B; D2-4) and the other three subregions were altered in the proximal half of the domain (Fig. 4A,C, P5-7). Each simulation was run for 24 hrs embryonic time and measurements were collected. In all simulations, neural crest cells populated most of the domain, with a cluster of cells at the lead of the stream (Fig. 4A-C). In the scenarios of increased domain growth distally, the model predicts that the furthest distance traveled by cells is reduced in comparison with the uniform growth model (U1 vs. D2-4, Fig. 4B). In contrast, when the proximal subregion of the domain grows twice as fast compared to the distal subregion (P5-7), the model predicts that cells travel further than with the uniform domain growth profile (Fig. 4C). When one subregion grows three times as fast as the other the probability that the neural crest stream breaks increases (example in T3, Fig. 4A). The furthest distance traveled by cells in each of the simulations (Fig. 4D). Together, the model simulations predict there are distinct differences in the distance traveled by neural crest cells when comparing proximal versus distal domain growth and suggest that the location of tissue growth is somewhat important to the neural crest cell migratory pattern. Also, there is a limit to the difference in growth rate between neighboring subregions beyond which neural crest stream breakage can occur.

**Figure 4:**
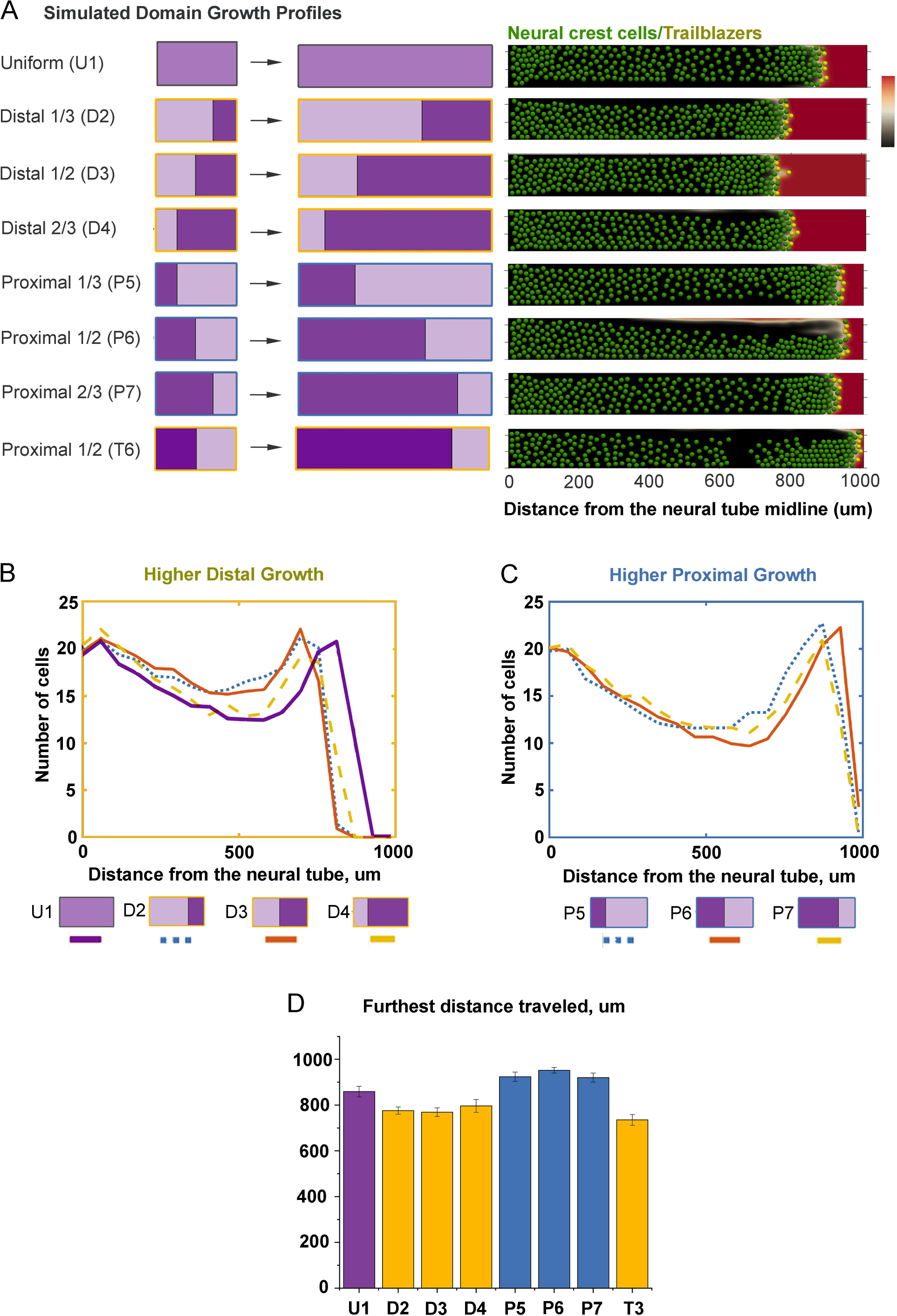
Modeling simulations of non-uniform tissue growth and the neural crest cell migratory pattern. (A) Schematics of tissue growth profiles. Darker color indicates a higher growth rate. The growth rate is doubled for models D2-4 and P5-7 and tripled the growth rate for T6. As the cells migrate into the domain they are advected at different rates and also contribute individual velocity to reach the end of the stream. Right, Example final distribution of cells (green) on domain with chemoattractant concentration (red/black color bar). Note the break in the stream in the simulation from model T3. Distribution of neural crest throughout the domain for each of the distal higher growth (B) and proximal higher growth (C). (D) Furthest distance traveled by cells under different growth profiles.

### Cell proliferation is heterogeneous in space and time within the head mesoderm and ectoderm along the dorsolateral neural crest cell migratory pathway

To determine the pattern of cell proliferation and growth within the head ectoderm and mesoderm, we performed pulsed BrdU labeling in ovo and measured the number of proliferating cells from confocal z-stacks in optically cleared chick embryos stages HH9-15 (Fig. 5A). We segmented nuclei based on methyl green staining, then categorized the tissue as either ectoderm (by distance from the surface), migrating neural crest cells (using HNK-1), or mesoderm (Fig. 5A,B). A cell was categorized as proliferating if the BrdU intensity was above a certain threshold (Fig. 5C). We find that overall, between HH9 and HH15, there is substantial inhomogeneity in cell proliferation in space and time (Fig 5C,E,F). For instance, at HH9, there are fewer proliferating cells near the midline and proliferation increases more laterally (Fig. 5C,F). By HH13-15 there are many subregions of high or low cell proliferation. We next specifically examined the percentage of BrdU-positive mesodermal cells along the dorsolateral neural crest cell migratory pathway from r4 to ba2 in 50 um bins starting at the midline to the end of the tissue (Fig. 5F). Before the neural crest cells exit the dorsal neural tube from mid-r3 to mid-r5 until HH13, the fraction of BrdU-positive mesodermal cells is highest distal to the invasive neural crest cells within a subregion approximately 100 um in length (Fig. 5F).

**Figure 5:**
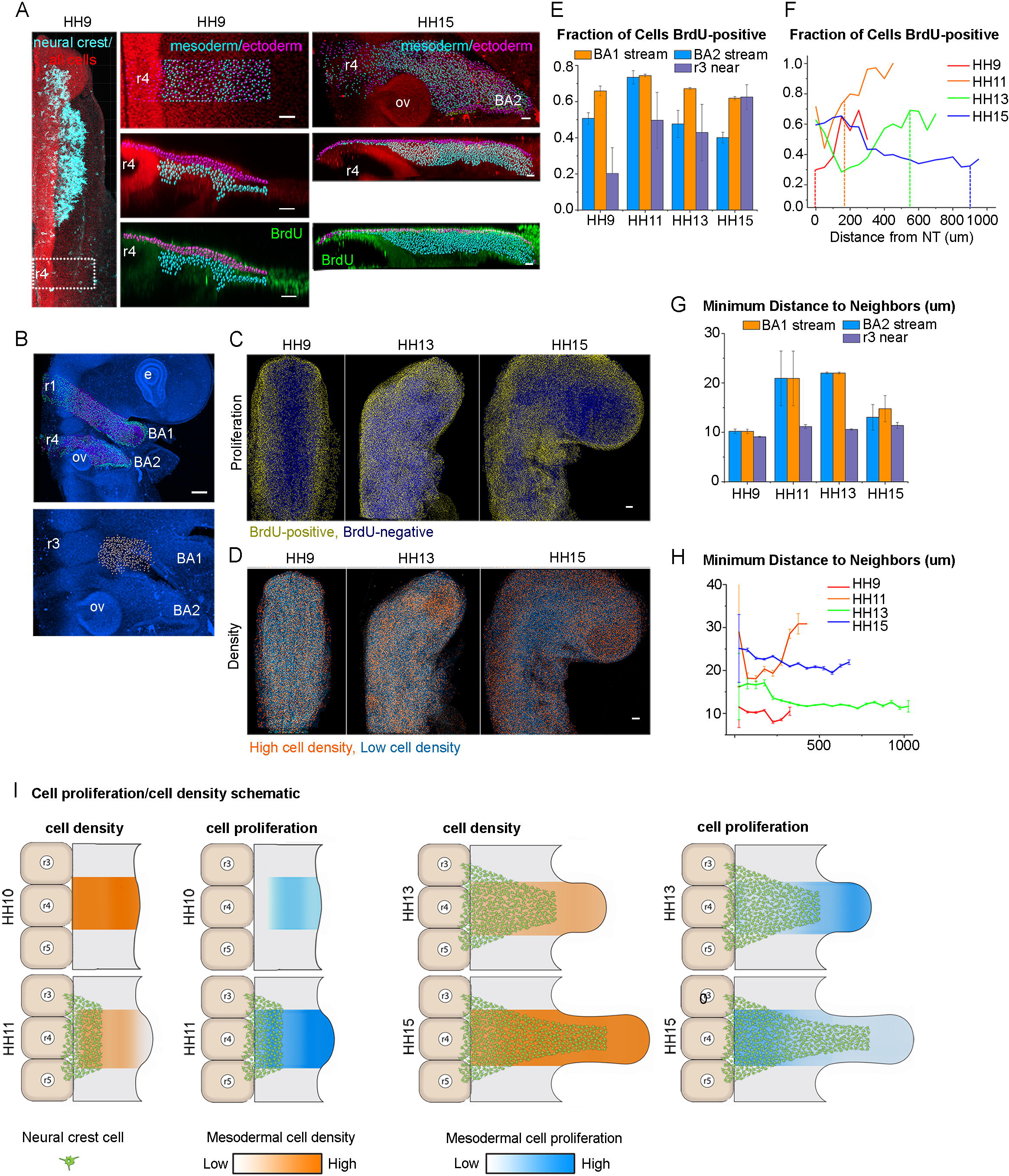
Proliferation and density of cells changes over time and throughout the tissue. (A) HH9 embryo with nuclei (red), neural crest (cyan) and BrdU (green). Second column of images shows area in dashed rectangle with spots segmented on the methyl green channel indicating location in the mesoderm (cyan) or ectoderm (magenta). Third column, HH15 labeled similarly. Scale bar 40um. (B) HH15 embryo. Mesodermal cells (cyan) and ectoderm (magenta). Below: area lateral to r3 studied marked with white spots. Scale bar 100um (C) Averaged images of embryos from HH09, HH13 and HH15 with yellow nuclei if the Brdu/methyl green intensity is in the top 40% of cells. (D) Averaged images of embryos from HH9, HH13 and HH15 with orange nuclei indicating the densest 40% of cells. Scale bar 100um. (E) Fraction of mesoderm cells in ba1, ba2 or adjacent to r3 domains that are positive for BrdU label in stages HH09-HH15 with SEM bars. (F) Fraction of cells BrdU-positive in 50um bins along the domain of r4 to the end of ba2. Dashed vertical lines represent the front of the neural crest stream. (G) Minimum distance to a neighbor averaged for all mesoderm cells for ba1 and ba2 streams, and area lateral to r3 with SEM bars. (H). Minimum distance between neighbors in 50um bins along the domain of r4 to the end of ba2. (I) Schematic of neural crest migration, tissue density (orange) and proliferation (blue) changes over time.

By HH15, the cell proliferation pattern changes to be higher in the paraxial mesoderm near the dorsal neural tube midline and uniform throughout the remainder of tissue towards ba2. Similar measurements were performed on the tissue moving towards branchial arch 1 (ba1) for comparison (Fig. 5B, Fig. S2). Along the neural crest cell migratory pathway towards ba1, the fraction of BrdU-positive cells was higher in the distal subregion at HH9, but this cell proliferation pattern becomes homogeneously distributed along the dorsolateral pathway from HH11-13 (Fig. S2). We measured a small increase in mesodermal cell proliferation in the paraxial mesoderm adjacent to rhombomere 1 (r1) at HH15 (Fig S2). For comparison, we also measured the fraction of mesodermal cells proliferating in the region distal to r3. Branchial arches 1 and 2 protrude outward from the embryo while this region does not grow as much (Fig. 5B, E). A smaller fraction of cells that proliferate in this area between stages HH9-HH13 but the fraction increases at HH15 to be more similar to the proliferation in ba1.

### Cell density remains constant within proliferating mesoderm implying tissue growth expansion

To address whether cell proliferation of the head mesoderm translated into an increase in local cell density and/or an increase in tissue size we measured the nearest neighbor distance between cell nuclei within the mesoderm between HH9 and 15 (Fig. 5D,G,H). We find that the subregions of high cell proliferation do not coincide with higher cell density (Fig. 5D, orange cells). When we examined the subregion neighboring the r4 neural crest cell migratory stream specifically, we found that the cell density is relatively constant through the subregion (Fig. 5H), with the exception of HH11. At HH9, mesodermal cells are tightly packed with an average of only 10 um, or a typical cell nucleus length, between nearest neighbors (Fig. 5G). As the tissue grows, mesodermal cells become less dense at HH11 and HH13 (24 um and 21 um on average, respectively) but pack in more tightly again by HH15 (13 um) (Fig. 5G). A schematic describing the neural crest stream development, proliferation and density can be found in Fig. 5I. These data support the concept that mesodermal cell proliferation gives rise to tissue growth. However, it is difficult to ascertain in what direction this growth takes place.

To address this, transgenic quail (typically approximately HH10) were mounted dorsal side down on EC culture for 405nm laser photoconversion of the Dendra2 photoconvertible protein (Fig. 6A,B). A region of interest was drawn roughly 100 um x100 umx50 um and scanned with the 405nm laser until the DendraRed signal no longer increased (Fig. 6A,B). Embryos were re-incubated for an additional 6-8 hours, imaged (Fig. 6A,B), and the dimensions of the photoconverted subregions were re-measured (Fig. 6C). We find that the photoconverted subregions expanded in the x-direction (along the dorsolateral neural crest migratory pathway and near the neural tube) but narrowed in the y-direction (parallel to the anteroposterior axis) (Fig. 6D). In the z-direction (dorsoventral), the photoconverted subregions changed a minimal amount (Fig. 6D). Together, these data suggest that cell proliferation within the paraxial mesoderm gives rise to growth of the tissue that is primarily along the proximal-to-distal axis in the direction of neural crest cell migration.

**Figure 6:**
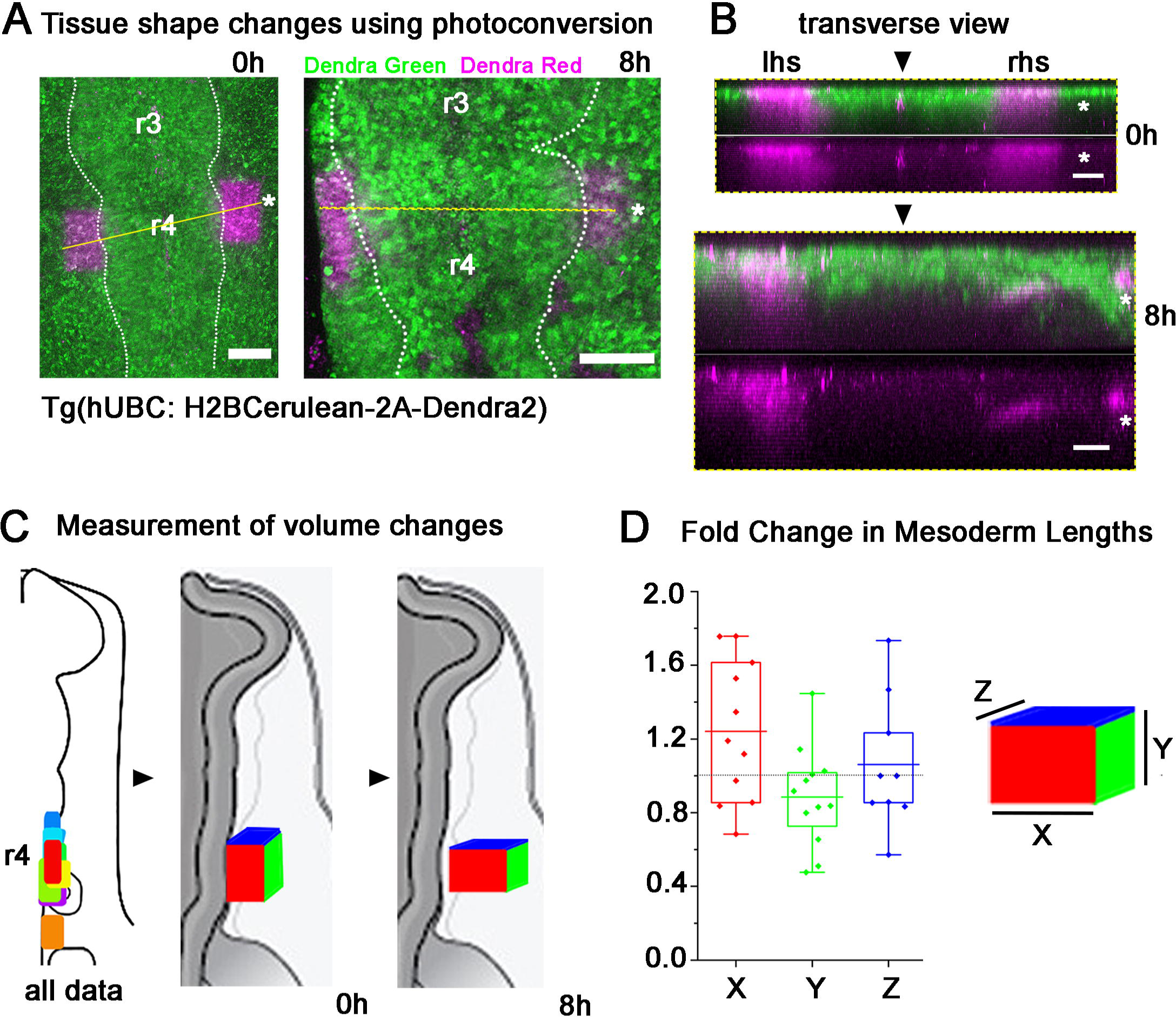
Photoconverted region of tissue grows more in the dorsolateral direction than anterior-posterior. (A) Rectangular regions where Dendra2 was photoconverted from green to magenta in HH9 embryo and imaged again 8 hours later. Scale bar 100um. (B) Transverse views through embryo in (A) at the dashed yellow line highlighting the depth of photoconversion. Scale bar 50um. (C) Schematic of embryos indicating locations of photoconverted regions at HH9. Also, schematic of embryo with box showing the growth of the tissue in the lateral (X) direction and compression in the anterior-posterior (Y) direction. Colors of the side of the box coordinate with colors in graph. (D) Fold change in size of mesoderm photoconverted region in X, Y and Z directions. No change marked with horizontal dashed line.

### The pattern of cell proliferation and tissue growth of the head mesoderm is altered after ablation of premigratory neural crest cells

In light of the cell proliferation and cell density measurements that showed tissue growth within the head mesoderm is distal to the invasive front of the neural crest cell migratory streams, we sought to better understand the influence of migrating neural crest cells on this pattern. To address this, we ablated the dorsal one-third of the neural tube in a bilateral manner from mid-r3 to mid-r5 in HH9 embryos to significantly reduce the number of premigratory neural crest cells (Fig. 7A). After 24 hrs of egg re-incubation to HH15, BrdU labeling was applied in ovo before embryos were harvested, fixed and processed for immunolabeling of the migrating neural crest cells with HNK-1. We measured the distal length from the neural tube midline to the brachial arches along the presumptive dorsolateral neural crest cell migratory pathway, and found this distance was significantly reduced in ablated embryos in comparison to the control (Fig. 7B). Further, of the premigratory neural crest cells that escaped ablation, we found these cells traveled a shorter distance towards ba2 (Fig. 7A,B). Examination of the pattern of BrdU-labeled mesodermal cells revealed that cell proliferation was higher along the length of the domain at every position but at the distal-most locations increased by 240% (Fig. 7C). Without the neural crest population binding to the VEGF expressed by the ectoderm, we examined the effects of increased VEGF on the mesoderm population by exposing mesoderm in culture to 100ug/mL VEGF. We found a significant increase in the percentage of dividing mesoderm cells (Fig. 7D,E). Thus, in the absence of migrating neural crest as in ablated embryos, increased proliferation of the mesoderm could be stimulated by increased exposure to VEGF.

**Figure 7.**
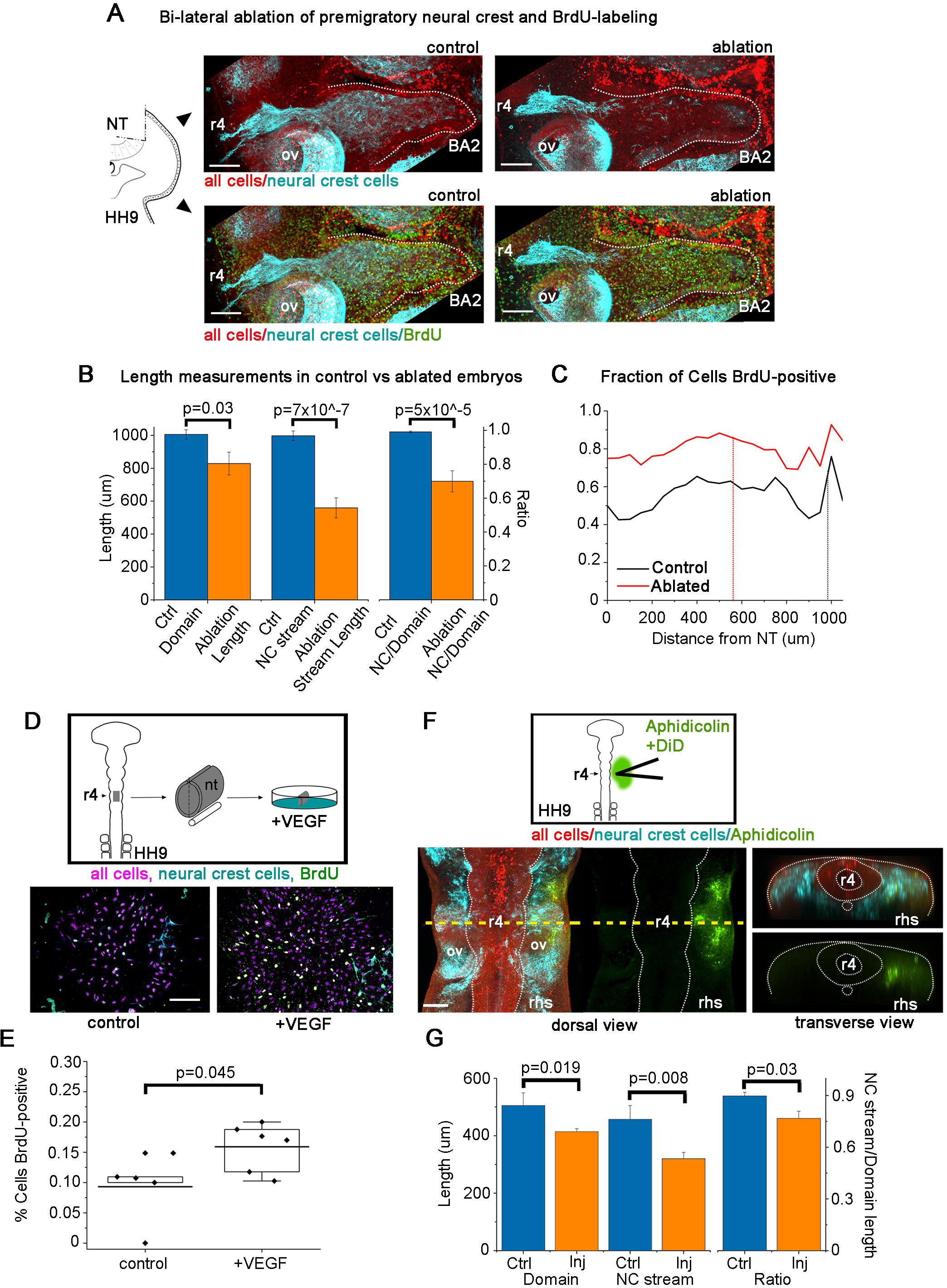
Physical ablation of premigratory neural crest cells or chemical inhibition of mesoderm growth show affects to both the neural crest cell migratory pattern and tissue growth. Representative r4 neural crest cell migratory stream 24 hrs post bi-lateral ablation of premigratory neural crest cells: (A) control (left) and ablated (right) stained with methyl green for all nuclei (red), HNK-1 for neural crest (cyan) and BrdU for proliferation (green). Scale bar 100um. (B) Length of domain and neural crest stream and the ratio for control and ablated embryos. (C) Fraction of non-neural crest cells in domain BrdU positive for control and ablated embryos. Vertical line indicates average length of neural crest stream. (D) Schematic and images of mesodermal cells in culture with or without exposure to VEGF for 16 hrs labeled with Hoescht for nuclei (magenta), HNK-1 for neural crest (cyan) and BrdU for proliferation (green). Scale bar is 100um. (E) Fraction of mesoderm cells in vitro that are BrdU-positive with or without VEGF. (F) Schematic and representative embryo injected with a cocktail of aphidicolin, hydroxyurea and DiD (green) after 8 hrs incubation. IHC performed with methyl green (red) and HNK-1 (cyan). Scale bar is 100um. Dorsal and transverse views of r4 axial level. (G) Length of domain and neural crest stream and ratio of treated and control embryos.

### Inhibition of mesodermal cell proliferation reduced domain size and length of the neural crest cell migratory stream

Computer model simulations presented above predicted significant changes in the neural crest cell migratory distance depending on the domain growth profile (Fig. 4). To slow the growth rate of tissue in the embryo, we microinjected a cocktail of Aphidicolin (1mg/mL), Hydroxyurea (5mM) and DiD into a broad subregion of the mesoderm and distal to the invasive front of the neural crest cell migratory stream in HH10 embryos (Fig. 7F). Aphidicolin and hydroxyurea are both well-known to inhibit the cell cycle and stop cell proliferation (Levenson and Hamlin, 1993; Mutomba and Wang, 1996). We microinjected control embryos with 10% DMSO and DiD. All microinjected embryos were re-incubated for 8 hrs then labeled with HNK1 to fluorescently label migrating neural crest and methyl green to mark all cells (Fig. 7F). We find that both the mesoderm within the dorsolateral domain and neural crest cell migratory stream were shorter in treated embryos (Fig. 7G). The ratio of the length of the neural crest cell migratory stream to the total length of the branchial arch tissue was significantly reduced (Fig. 7G). Thus, by chemically inhibiting cell proliferation in the mesoderm, we were able to reduce tissue growth and alter the pattern of neural crest cell migration.

## DISCUSSION

Previous static studies have suggested that head mesoderm (through which neural crest cells travel) expands in a uniform manner in space (McLennan et al., 2012) and neural crest cells are passively carried along by the mesoderm (Noden and Trainor, 2005; Evans and Noden, 2006). Our discovery of the inherent motility of individual mesodermal cells in culture, and dramatic changes in mesodermal cell behaviors in the presence of cranial neural crest cells or ectoderm, suggested a more complex and dynamic choreography of vertebrate head morphogenesis. By integrating mathematical modeling with in vivo imaging and experiment, we were able to predict changes in the neural crest cell migratory pattern under different (non-uniform in space) mesoderm growth profiles and compare this to measured changes in mesodermal cell proliferation, cell density, and cell trajectories and to experimental manipulations of migrating neural crest cells and mesodermal cell proliferation. Our results provide a detailed picture of the rich spatio-temporal pattern of ectoderm and mesodermal cell dynamics and cell-cell interactions with the migrating neural crest cells.

Head mesodermal cells in culture displayed inherent dynamic migratory behaviors in vitro, suggesting these cells are capable of active cell movements in vivo and exchange of neighbor relationships. Mesodermal cells isolated from the chick head paraxial mesoderm adjacent to the rostral hindbrain dispersed as individuals throughout the culture dish in a similar manner to neural crest cells that exit cultured neural tube explants (Fig. 1). Mesodermal cells in culture moved 81% slower but 155% more directed than neural crest cells in vitro (Fig. 1). This was surprising since paraxial and lateral mesoderm have previously been assumed as passive tissue (Noden and Trainor, 2005; Evans and Noden, 2006). Interestingly, when placed in co-culture with neural crest cells, both mesodermal and neural crest cells migrated faster and freely intermingled (Fig. 1). Thus, the inherent motility of head mesodermal cells and dynamic interplay with the neural crest and ectoderm in culture mean that cell behaviors must be closely coordinated in vivo to create the stereotypical pattern of the head and branchial arches.

Ectoderm and mesodermal cells displayed large scale collective whirling motions prior to neural crest cell emergence. Whirling patterns of cells have a rich history from observations of confluent fibroblast cultures (Elsdale and Wasoff, 1976) and the insect integument (Nubler-Jung, 1987), and are borne out from mathematical models based on the mechanochemical properties of tissue (Oster et al., 1983). More recently, whirling motions have been visualized during vertebrate gastrulation in the tailbud of zebrafish and chick embryos, especially within the mesodermal progenitor zone (MPZ) (Zamir et al., 2006; Lawton et al., 2013; Mongera et al., 2018). Curiously, the observance of a large scale, circular structure has previously been reported in chick head mesoderm and visualized using stereo scanning electron microscopy (Meier, 1981; Jacobson, 1988). They reported paired mesodermal cell disks which they termed ‘somitomeres’ with a diameter ranging from 135 to 240 um, appearing in an anterior-to-posterior order starting at about HH4 through HH9. Cells were arranged around one or two central cells with a tuft of processes pointing toward the ectoderm (Meier, 1981; Jacobson, 1988). When we measured ectoderm and mesodermal cell movements within presumed somitomeres, we found that cells traveled in a counter-clockwise direction on the right-hand-side of the embryo (and clockwise on left-hand-side) at a speed of 0.2 degrees per minute (Fig. 2 C-E). Further analysis of mesodermal cell-cell interactions and changes in tissue boundary geometries or molecular signals in the microenvironment in the vertebrate head should help us better understand the functional relationship between whirling motions, tissue growth, and the emergence of the migrating neural crest cells.

Mesodermal cells rapidly transitioned to directed motion towards the branchial arch target sites and interacted with invading neural crest cells, suggesting a close coordination between the neural crest and mesodermal cells to expand tissue growth and morphogenesis of the periphery. Mesodermal cells that interacted with neural crest cells displayed more directed cell trajectories (Fig. 1, Fig. 3F, circles). Faster moving neural crest cells passed through slower moving mesodermal cells (Fig. 3F, squares) and some mesodermal cells were displaced either lateral or off the migratory pathway after interacting with a neural crest cell (Fig. 3F, triangle). The fact that neural crest and mesodermal cells interacted was not surprising given previous data from quail-chick chimeras or viral labeling studies that reported populations of these two cell types developed in close registration (summarized in (Le Douarin and Kalcheim, 1999); reviewed in (Noden and Trainor, 2005)). However, it was stunning to observe the rich individual mesodermal and neural crest cell-cell interactions and rapid movements of neural crest cells to overtake individual mesodermal cells en route to the branchial arches (see Movie 8). We did not observe neural crest cells to be passively carried along by the underlying moving mesodermal tissue (Evans and Noden, 2006). Intriguingly, neural crest cells that were observed to displace loosely connected mesodermal cells out of the way would require cell contact with enough force to change the direction of the contacted cell. Further time-lapse studies of local mesodermal and neural crest cell-cell interactions combined with an examination of the expression of cell receptor-ligand pairs should help to better understand the dynamic relationship between these two cell types and the transition of the mesodermal cells to lateral movement.

Computational modeling allowed us to tackle the question of how different growth profiles affect the neural crest cell migratory pattern. When we tested six profiles simulating non-uniform mesoderm growth (three out of six profiles consisting of faster growth in the proximal subregion and the other three profiles with faster growth in the distal subregion), we found that the neural crest cell migratory distance traveled was reduced in the presence of faster distal growth (Fig. 4A,B,D). This appeared to be due primarily to reduced cell advection on the proximal subregion. In contrast, the model predicted neural crest cells traveled further along the domain when the proximal domain grew faster, as would be expected (Fig. 4A,C,D). This appears to be due primarily to neural crest cell advection through the faster growth domain. Only with a large difference between growth rates of the neighboring subregions do we see neural crest stream breakage (Fig. 4A). While the neural crest stream length changes by as much as 20% as we vary the growth profile along the domain, the formation of a continuous stream is reliable in the model. Also, since stream formation still occurs during embryo manipulations (ablation or low proliferation rate) this would imply a robustness in the ability of neural crest to form streams of cells under a variety of growth conditions.

The combination of cell proliferation and cell density measurements of the chick head mesoderm showed that the tissue growth is non-uniform in space, suggesting more complex cell dynamics than previously thought. Cell density remained constant within proliferating cell mesoderm implying tissue growth expansion. Further, photoconversion of small tissue volumes as well as time-lapse imaging showed that the growth of the mesoderm was preferential in the proximal-to-distal direction (Fig. 6D). Using a simple length measurement along the surface ectoderm from the dorsal neural tube to the forming ba2 revealed logistic growth during the developmental stages from prior to neural crest cell emergence through cell invasion of the branchial arches (McLennan et al., 2012). Previously, we simply assumed uniform growth spatially within the tissue (McLennan et al., 2017) in the absence of detailed analysis. However, our present careful cell proliferation analysis above showed spatially non-uniform tissue growth. Interestingly, these data support our observation that neural crest cells establish cell polarity after exiting the dorsal neural tube within a subregion approximately 150um from the dorsal midline and change gene expression from an epithelial-to-mesenchymal to directed cell migration signature after encountering signals within the paraxial mesoderm (Morrison et al., 2017). With the mesodermal tissue growth then shifting to distal to (or ahead of) the invasive front, neural crest cells must then actively migrate to stay in pace with the branchial arches (Fig. 5F). In analogy, the neural crest cells are on a moving walkway in which the different sections of the walkway move faster/slower over time. How the neural crest cells modulate advection and active migration while traveling through the mesoderm will be exciting to examine in the future.

There were contradictions between the model simulations of non-uniform domain growth and our experimental results. First, we measured higher distal growth rates in the mesodermal tissue during neural crest cell migration. However, the model predicted that neural crest cells traveled farther with proximal growth conditions (Fig. 5F vs. Fig. 4A,D). This could be explained by the limitation that we did not include a reduction in tissue growth in the model that was measured experimentally during the progression from HH13-to-HH15 (Fig. 5). If the most distal subregion of the domain halts proliferation towards the end of the simulation, then the neural crest would be able to “catch up” to the end of the tissue, resulting in a ∼1000 um continuous neural crest stream to the end of the branchial arch. Second, we observed an approximately constant speed of neural crest cells for at least the first 300 um of invasion (Movies 7,8). However, the model predicted that neural crest cell speed increased the further a cell was from the neural tube (because of growth-driven advection upon observed cell speed). It is challenging to separate the impact of passive growth-driven advection from active neural crest cell motility in our time-lapse imaging data. In the absence of such data, it is not possible to tell whether cells modulate their speed during migration. Neural crest cells have been observed to momentarily slow during a transition from migrating to dividing but quickly speed up to regain their position in the stream (Ridenour et al., 2014). Thus, there appears to be a subtle balance between advection and independent motion of the migrating neural crest cells.

The pattern of cell proliferation and tissue growth of the head mesoderm was altered after ablation of premigratory neural crest cells, suggesting VEGF (or other growth factor signals) acts on both neural crest migration and mesoderm proliferation simultaneously. After ablation of premigratory r4 neural crest cells, ba2 forms but is smaller than in control embryos (Fig. 7A,C). Further, we learned that cultured mesodermal cells overproliferated in response to VEGF added to the media (Fig. 7D,E). Thus, these data support previous observations that branchial arch tissue can form independent of the neural crest cells because of the neural crest-independent signaling (Veitch et al., 1999), and the increase in proliferation of mesodermal cells may result from an overabundance of VEGF. However, the downstream morphogenesis of muscle and bone would still be affected (Noden, 1988; Veitch et al., 1999).

We also asked the converse question, if the branchial arch were to diminish in size, what effect would this have on the migratory neural crest stream? Chemical inhibition of cell proliferation in the mesoderm led to reduced length of ba2, but the neural crest cell migratory stream length was also significantly reduced (Fig. 7G). We anticipated that the neural crest cells would travel further into the branchial arch with a shorter domain, however since the neural crest cells traveled a shorter distance through the migratory domain, this implied a signaling interplay between the independent motion of the neural crest and growth activity of the mesoderm that will be interesting to explore in the future.

In conclusion, our results demonstrate that the neural crest cell migratory domain grows in a non-uniform manner in space and time during vertebrate head morphogenesis. Paraxial mesodermal cells are inherently motile and display collective cell movements that transition from large-scale whirling (prior to neural crest cell emergence) to directed trajectories towards the branchial arches. By simultaneously observing ectodermal, mesodermal and neural crest cell behaviors, we have unraveled the complex cell dynamics that lead to head and branchial arch morphogenesis. By testing a number of hypothetical non-uniform in space tissue growth scenarios in silico, we arrived at a better understanding of what would be possible in order to support neural crest cell migratory stream patterning. Empirical measurements then revealed spatiotemporal changes in tissue growth expansion that begin within the paraxial mesoderm and transition to the distal subregion of the forming branchial arches over time. These data support a model wherein neural crest cells must balance active motility and growth-driven advection to preserve movement as a collective cell population. Our manipulation of the neural crest cell migratory domain by ablation of the premigratory neural crest or inhibition of mesodermal cell proliferation strengthen the idea that interactions between the two cell populations stimulate tissue growth. Lastly, our detailed analysis of avian head morphogenesis and neural crest cell migration provide a framework to integrate single cell genomic data and comparisons between different vertebrate research organisms.

## MATERIALS AND METHODS

### Embryos and in ovo labeling

Fertilized white leghorn chicken eggs (Centurion Poultry) or Tg(hUBC: H2BCerulean-2A-Dendra2) quail (Translational Imaging Center, University of Southern California) were incubated at 38° C in a humidified incubator until the desired Hamburger and Hamilton (HH) stage (Hamburger and Hamilton, 1951) of development. A 0.3M sucrose solution was mixed 1:1 with DiD vibrant dye solution (V22887 Thermo Fisher) and injected into the neural tubes of embryos at HH9 for neural crest labeling.

### In vitro assays

For mesoderm and neural tube cultures, glass bottomed petri dishes (P35G-1.5-20C, Mattek) were coated with 20ug/ml of fibronectin (F1141; Millipore Sigma) and 20ug/ml of poly-D-lysine (P7886; Millipore Sigma) for 30 minutes then allowed to dry. Lateral mesoderm tissue from the r4 axial level was isolated from stage HH6-8 chick embryos by creating transverse cuts at the r3/r4 and r4/r5 borders and then removing neural tube, ectoderm and endoderm with a sharpened tungsten needle and forceps. The remaining mesoderm was placed onto prepared glass bottomed dishes. Isolated neural tube explants were prepared similarly to (McLennan et al., 2010) and placed in a separate dish or near the mesoderm tissue. Cells were cultured in Ham’s F-12 nutrient mix (11765047; Thermo Fisher) supplemented with Pen/Strep and B27 (17504044, Thermo Fisher) at 37° either in a tissue culture incubator or in an environmental chamber on the microscope. For mesoderm experiments with VEGF, 100ug/mL of VEGF (293-VE-010, R&D systems) was added to the F-12 media for 16-hour incubation or PBS for control. Cells were labeled with 5ug/mL of Hoechst 33342 (b2261, Millipore Sigma) for 10 minutes either for live imaging or after immunohistochemistry. Glass bottomed dishes for in vitro cultures were placed in a 6-well microscope stage insert inside a heated environmental chamber on an LSM-800 confocal microscope (Zeiss) and imaged every 5 minutes, using a Plan Apochromat 10× 0.45 NA M27 objective, single z plane, with a minimum of 405 laser intensity for 16 or more hours. Analysis was performed on many cells from multiple time lapses (3 mesoderm/neural tube, 5 mesoderm only, 4 neural tube only, 3 mesoderm and ectoderm). Significance was determined with 2-tailed Student’s T-test throughout the manuscript.

### Antibody labeling

For proliferation studies in vivo, the vitelline membrane was removed from the head of the embryo then 50uL of BrdU labeling reagent (000103, Thermo Fisher) was dropped on top. The embryos were reincubated for 30 min then fixed in 4% paraformaldehyde overnight at 4°. The head and first four somites were isolated from the trunk then embryos HH13 and older were bisected down the midline. For immunohistochemistry, fixed embryos were incubated for 10 minutes in 1N HCl on ice, 10 minutes in 2N HCl at room temperature followed by 20 minutes at 37°. After 12 minutes in Borate buffer, the embryos were incubated with PBS+1% TritonX and finally 10% goat serum (16210072, Thermo Fisher) block. In vitro cultures were prepared by adding 1% BrdU labeling reagent and cultured for 1 hour before fixing with 4% paraformaldehyde for 30 minutes at room temperature. In vitro protocols only require 15 minutes 1N HCl at 37° and 15’ Borate buffer before washing and blocking. Embryos or cells were labeled with primary antibodies BrdU (1:200, pa5-32256, ThermoFisher, Lot TH2627941A) and HNK-1 (1:25, TIB-200, ATCC) in block overnight at 4°. After extensive washing in block at room temperature, secondary antibodies goat anti-rabbit IgG Alexa Fluor 488 and goat anti-mouse IgM AlexaFluor 546 (A-11008 and A-21045, Thermo Fisher) were applied at a 1:500 dilution overnight at 4°. For a nuclear stain, Methyl Green (1:5000, M8884 Millipore Sigma, (Prieto et al., 2015)) was added to embryos after immunohistochemistry for several hours. Embryos with no primary antibody were used as control. Embryos were cleared for imaging using 80% FRUIT clearing buffer (Hou et al., 2015) overnight at 4° and mounted for imaging in 80% FRUIT buffer on glass slides. At least 3 and up to 5 embryos’ data were combined for each stage.

### Embryo Manipulations

Bilateral ablations of the dorsal 1/3 of the neural tube were performed on HH9 embryos in ovo. Using glass needle and forceps, a cut was created down the midline from r3 to r5 and the dorsal aspect neural tube was removed. Control embryos were opened, staged and re-sealed. After 24-hour incubation, embryos were treated with BrdU reagent as above for 30 minutes then harvested and processed for immunohistochemistry. Eight embryos control and treated were analyzed. To halt proliferation in the mesoderm, Stage HH10 embryos in ovo were injected with a freshly prepared cocktail of 100ug/mL Aphidicolin (A0781, Millipore Sigma), 5mM hydroxyurea (H8627 Millipore Sigma), and 10% DiD. Injections were made at the axial level of r3 to r5 as lateral as could be reached but still in the mesoderm. DiD was used to visualize the area injected and give a rough estimate of where the chemicals distributed. Control embryos were injected with 10% DMSO and DiD. After injection, embryos were resealed and incubated for 8 hours then fixed and processed for immunohistochemistry. Thirteen injected and eight control embryos were used for analysis.

### Microscopy

Live embryos for time lapse imaging were mounted on EC culture dorsal-side down beginning at HH9 or 10 modifying the protocol of (Chapman et al., 2001; McKinney et al., 2016) such that the EC culture was plated with only 500uL of liquid to reduce light scattering. Confocal z-stacks using 488nm laser to image Dendra2 of the transgenic quail and 633 for DiD were collected every 7 minutes for up to 12 hours with a 10× 0.45 objective. Embryos used for photoconversion were mounted similarly on EC culture. Regions approximately 100×100 um for Dendra2 photoconversion were chosen on both sides of the embryo if possible. 5% 405nm laser was scanned in this region and fluorescence intensity of Green Dendra and Red Dendra were monitored in a second channel until the Red intensity no longer increased. A larger, tiled z-stack image of the whole head was then acquired. Embryos were transferred to an incubator for 6 to 8 hours then removed from the culture dish by the paper ring, mounted on a glass slide and immediately re-imaged in a tiled z-stack. Fixed embryos were imaged in 80% FRUIT clearing buffer in tiled, z-stack images with a Plan Apochromat 10× 0.45 NA objective on an LSM-800 microscope. Microscopy was optimized for imaging AlexaFluor 488, AlexaFluor 546 and methyl green.

### Image Analysis

Both in vitro and in vivo time-lapse imaging series were concatenated and aligned (Preibisch et al., 2010) in Fiji (Schindelin et al., 2012). Post alignment, images were imported into Imaris (Bitplane AG) for tracking cells with fluorescent label. In vitro mesoderm and neural crest were tracked by hand due to the single color labeling of the nuclei. Embryonic cells were tracked semi-automatically with Imaris. Track position and statistics were exported to MATLAB (Mathworks Inc) scripts for further analysis. Distance between random cells (Fig. 2I,3E) was calculated only for the time both cells existed. If a division occurred, one random daughter was picked. Images of BrdU, methyl green and HNK1 labeled embryos were first stitched in Fiji (Preibisch et al., 2009), then imported to Imaris for segmentation. Images were smoothed and attenuation corrected in Imaris then the domain around the r4 neural crest stream was isolated by hand. For younger embryos with little or no neural crest present, an area was selected dorsal to the neural tube at the r4 axial level across to the lateral end of the tissue directly across from r4. The width was chosen based on shape of the neural tube. For older embryos, the region of interest was selected from around the neural crest stream down to the most lateral end of the tissue. Similar procedures were followed for the stream of neural crest targeting ba1. The area lateral to r3 was isolated by creating a 100 um circle from the ectoderm between the ba1 and ba2 streams (including the ectoderm). Segmentation was performed on the methyl green signal. Separation of the ectoderm was achieved using a distance transformation to the original domain region and some hand-editing. While the HNK-1 signal is a membrane label, the channel is bright enough to create an intensity cutoff for neural crest identification at the nucleus. The bright BrdU signal allowed us to create an intensity cutoff for proliferation. The average intensity, position and minimum distance to a neighbor were exported to MATLAB scripts for calculating the fraction of cells proliferating and minimum distance to a neighbor in 50 um bins down the length of the domain. MATLAB scripts can be obtained through Stowers Original Data Repository upon publication.

### Computer Model Simulations

We assume that there are two types of cells, namely “leaders” and “followers”. There are a fixed number of leaders who undertake a biased random walk with volume exclusion up a cell-induced gradient of chemoattractant. The leaders then perform a biased random walk by extending three filopodia in random directions per time step. These filopodia are assumed able to sense the concentration of chemoattractant at their tip and the cell then moves in the direction of the highest concentration sensed, provided it is higher than that at the position of the center of a cell. If this is not the case, then the cell moves in a random direction. On the other hand, followers are either in a chain or move randomly. A chain consists of a group of followers that are close to each other and at least one of the cells is close to a leader. All the followers in a chain move in the same direction as the leader that is at the front of the chain. If a follower is relatively close to more than one leader, it may join a few different chains, then the cell randomly chooses which one to follow. We include a simplified version of phenotype switching between leaders and followers based on the position of a cell within a migratory stream. We use experimental data on the initial and final length of the domain. We consider seven different types of domain growth (Fig. 4). Initially, we consider uniform domain growth, then we split the domain in two parts, which we call proximal and distal subregions (Fig. 4). We carry out three different studies: different percentages of the total length undergo faster proximal growth or distal growth (Fig. 4A; D2-4 and P5-7), and one in which the difference in the elongation between the two parts is a factor of three (Fig. 4A; T3). We assume exponential growth in time and pick the growth rates in such a way that each of the growth profiles leads to the same total domain length after 24 hours. We run twenty simulations for each model and store statistics after 24 hours on the density of cells along the domain, the furthest distance travelled (which we quantify as the average distance travelled by the leader cells), and the fraction of the simulations with more than 20% follower cells not in chains (a measure of the likelihood of stream breakage).

## ACKNOWLEDGEMENTS

PMK would like to acknowledge the kind and generous funding from the Stowers Institute for Medical Research. We would also like to thank David Huss, Rusty Lansford, and the Translational Imaging Center at the University of Southern California for use of the transgenic quail. In addition, we thank Dave Lei and Ashley Young for contributions to imaging and image analysis as part of the Stowers Summer Scholars Program.

## DATA AVAILABILITY

Original data underlying this manuscript can be accessed from the Stowers Original Data Repository at https://www.stowers.org/research/publications/odr

---

**Supplementary Figure 1. Mesoderm explants rarely contain neural crest cells.** (A) An example neural tube explant labeled with Hoechst and HNK1 by IHC with many labeled neural crest migrating away from the neural tube. (B) a mesoderm explant labeled similarly with only a few neural crest.

**Supplementary Figure 2. Proliferation of mesoderm cells in between r1 and ba1.** Fraction of cells BrdU positive in 50um bins along the domain between r1 to the end of ba1 (domain highlighted in Figure 5B).

**Supplementary Movie 1. Lateral mesoderm cells in culture migrates dynamically**. Hoechst labeled explanted lateral mesoderm from HH10 chick embryo imaged in 5-minute intervals for approximately 16 hours.

**Supplementary Movie 2. Neural tube culture illustrating the dynamically migrating neural crest**. Hoechst labeled neural tube explant imaged in 5-minute intervals for approximately 16 hours. Many neural crest can be seen exiting the neural tube and exploring the dish.

**Supplementary Movie 3.Neural crest cells interact with mesoderm cells in vitro**. Hoechst labeled neural tube explant and mesoderm explant imaged in 5-minute intervals for approximately 16 hours.

**Supplementary Movie 4. Mesoderm adheres to ectoderm when cultured together.** Hoechst labeled and mesoderm and surface ectoderm explant imaged in 5-minute intervals for approximately 16 hours

**Supplementary Movie 5. Gross morphological changes during neural crest migration at the r4 axial level**. Transverse view of the r4 axial level of a whole transgenic quail embryo culture (green) with DiD labeled neural crest and neural tube (magenta). The tissue bends and grows ventrally while cells stream laterally. Confocal z-stack images taken in 7-minute intervals for several hours.

**Supplementary Movie 6. Whirling motion of ectoderm and mesoderm can be observed lateral to r3**. Dorsal view of transgenic quail (green) with DiD labeled neural tube and migratory neural crest cells (magenta). Before the neural crest emerge from the r4 axial level a counterclockwise rotation of cells can be observed at r3. Confocal z-stack images taken in 7-minute intervals for several hours.

**Supplementary Movie 7. Neural crest migrate much faster than the ectoderm sliding under the tissue.** Transverse (top) and Dorsal (bottom) views of transgenic embryo (green) with DiD labeled neural tube and neural crest (magenta) of HH12 stage quail embryo. The cyan marked ectoderm cells move very little compared to the neural crest marked in yellow that slide beneath. Images taken in 4.5-minute intervals for several hours.

**Supplementary Movie 8. neural crest bulldoze through the mesoderm displacing cells**. Dorsal view of transgenic quail embryo (green) with DiD neural crest (magenta). Three mesoderm cells are highlighted in white and some of the neural crest they interact with are highlighted in magenta. The mesoderm cells can be seen to passed by the neural crest and diverted posteriorly (1, squares), become more directed and seem to be pushed (2, circles), or move far posterior to its original location and eventually passed by the neural crest stream (3, triangle).

